# Human Autosomal Nucleotide Positions Differing from Bonobo Instead Match Pig

**DOI:** 10.1101/2024.08.14.607926

**Authors:** Eugene M. McCarthy

## Abstract

**Overview:** To examine the hybrid hypothesis of human origins, a novel data mining program, BOOMSTICK, was used to scan the euchromatic portions of two target genomes, those of *Homo sapiens* and *Pan paniscus*. Each of the two genomes were broken up into 100-kB segments, each of which was searched for matches to a large set of porcine queries. All scans sought matches to the same set of 813,194 40-mer nucleotide queries randomly selected from the genome of *Sus scrofa* (domestic pig). For each of the two study organisms, mean segmental match rates (MSMRs) were then calculated for all segments in each of three categories: those segments occurring on autosomes, those on the X chromosome, and those on the Y chromosome.

**Results:** In scans of single-copy regions (euchromatin) in both their Y chromosomes and their autosomes, it was found that the number of matches to randomly selected porcine queries was higher in humans than in bonobos. When autosomes were compared, matches were 1.3% higher in humans than in bonobos. This figure is equal to the percentage of human autosomal nucleotide positions bearing nucleotides that match in pig but not in bonobo. Remarkably, it agrees with the percentage of autosomal nucleotides previously reported to differ in bonobos and humans. So, the results of this study indicate that essentially all the nucleotide positions that differ in humans and bonobos, are the same in humans and pigs. In addition, the number of matches to pig queries found on the human Y chromosome was 34.5% higher than on the bonobo Y, and 12.4% higher than on the chimpanzee Y (the chimpanzee figure may be the more reliable of the two, since the bonobo Y nucleotide sequence file scanned contained only unlocalized scaffolds). MSMRs for the human and bonobo X chromosomes did not significantly differ.

## Introduction

Many researchers now deem the chimpanzee (*Pan troglodytes*) and the bonobo (*Pan paniscus*), to be the closest living relatives of *Homo sapiens*, which is to say that they think humans and panins (i.e., apes of the genus *Pan*) share a more recent common ancestor than does either with any other kind of organism (McNulty 2016). Under this view, two lineages, one leading to modern humans and one to the extant panins, gradually diverged from this common ancestor. This divergence is pictured as a steady accumulation, in each of the two lines, of the traits that today distinguish *Homo* and *Pan*. Here, this widely accepted belief will be termed the *divergence hypothesis*.

An alternative notion, what will here be called the *hybrid hypothesis* of human origins, is the idea that the human line found its origin in hybridization between a panin and some other kind of animal. Hybridization is the production of composite offspring through the interbreeding of distinct kinds of organisms. Such offspring, and their descendants, are termed hybrids. Hybridization has long been recognized as an important factor in the creation of new kinds of plants. But for most of what was called the “modern synthesis” era, biologists generally argued that it played no significant role among animals (Taylor and Larson 2019). In recent years, however, a consensus has been forming that hybridization can, in fact, play a generative role in animal evolution (ibid.). Hybridization can create new combinations of preexisting alleles (Marques et al. 2019) and/or chromosomes (McCarthy et al. 1995). Even if most such new combinations were deleterious, or even lethal, some might well be beneficial. Such new combinations could, at least theoretically, result in reproductive isolation between hybrid populations and the parental populations that crossed to produce them (McCarthy et al. 1995; Marques et al. 2019; see also Grant and Grant 2009). It has been proposed that such events may trigger explosive adaptive radiations (Gillespie et al. 2020; Marques et al. 2019; McGee et al. 2020; Meier et al. 2017). McCarthy (2008a) went as far as to argue that hybridization is likely the primary force in the creation of new forms of life.

Here reported are the results of a large-scale study evaluating the question of whether the genome of *Homo sapiens* might contain important contributions from organisms outside what is usually considered its immediate evolutionary clade. The motivation for this study will be better understood if the reader first considers how biologists identify hybrids.

When an organism is produced under controlled conditions and its parents are known, it is also known whether it is a hybrid. But the parentage of creatures found in a natural setting can only be deduced (McCarthy 2006). Under such conditions, if a biologist wishes to know whether a given individual had a hybrid origin, the initial evaluation will be phenotypic. Hybrids usually mix the traits of their parents and are intermediate with respect to most physical measurements. Say an ornithologist finds a bird that mixes the traits of a Baltimore Oriole (*Icterus galbula*) with those of a Bullock’s Oriole (*I. bullockii*). Suppose, too, that it’s intermediate in most respects between those two birds when its various parts are measured. She will conclude that it’s a hybrid of the two. In many cases, such an analysis of traits is the only evidence adduced when a scientist claims that a given organism is a hybrid. In some cases, as in the present one, such findings will prompt subsequent genetic studies.

McCarthy (2008b) employed the phenotypic method in an initial evaluation of the question of whether humans might be of hybrid origin. A search of the literature revealed that humans seemed to mix the traits of panins with those of pigs, and that they are intermediate between panins and pigs with respect to many physical traits. In total, the study documented about a hundred such traits that distinguish humans from panins. If many such traits were not found in pigs, then those that were could reasonably be attributed to convergent evolution. Such an explanation, however, seems tenuous given that nearly every such trait that McCarthy identified was also found in pigs. This strong tendency for such traits to be found in pigs supports the notion that they are instead relics of ancient hybridization, specifically, hybridization between panins and pigs. Moreover, detailed phenotypic analysis pointed to *Pan paniscus* as the specific panin that participated in the cross. This conclusion agrees, too, with the fact that bonobos share slightly more nucleotide identities with humans than do chimpanzees.

Thus, the hybrid hypothesis can be phrased more precisely as the idea that the human line had its origin in prehistoric hybridization between pigs and a particular panin, the bonobo. The present study is intended as an initial investigation of whether genetic evidence is also consistent with this hypothesis.

An immediate and obvious objection to the idea that humans are pig-ape hybrids is the fact that such a cross would be interordinal. The infeasibility of such crosses, though not well supported by observation, has become one of the unquestioned axioms of biology. If it could in fact be established that interordinal hybrids cannot occur, then the hybrid hypothesis could be rejected at once. But to address this issue, McCarthy collected information bearing on the question of the limits of hybridization among animals. The resulting publication (McCarthy 2021) lists about fifteen hundred separate reported cases involving interordinal animal hybrids, many of which were allegedly produced without human intervention. The existence of these many reports shows that interordinal hybrids can in fact be produced, sometimes even in a natural setting. And, though such hybrids may be rare measured on a human timescale, measured on an evolutionary one they are extremely common. Such events therefore should not be dismissed, especially given that they could produce large effects. In the present case, their documented existence also makes the idea more plausible that bonobos and pigs might have prehistorically crossed.

Physically, humans look more like bonobos than pigs, and they share more nucleotide identities with the former than with the latter. Therefore, under the hybrid hypothesis, one would assume that any initial hybridization producing a first (F_1_) generation was followed by one or more generations of backcrossing to bonobos. A first backcross (B_1_) generation would occur when an F_1_ hybrid produced offspring with a bonobo, a second backcross (B_2_) generation, when a B_1_ hybrid went on to produce offspring with a bonobo, and so forth. In a cross between pig and bonobo, the F_1_ hybrids would be expected to have about half their genetic material from pig and half from bonobo (but not exactly half, since the bonobo and pig genomes differ in size). After one backcross, the B_1_ genome would be about one-quarter pig and three-quarters bonobo. However, due to variation during meiosis in the gonads of F_1_ hybrids, resulting from both differences in recombination (during metaphase I) and in segregation (during anaphase I and II), the relative amounts of pig and bonobo DNA present in the genome would vary widely from one backcross individual to another. There would be genetic and phenotypic variation, too, among the second backcross hybrids. But on average, a B_2_ genome would be about one-eighth pig and seven-eighths bonobo. Another backcross would reduce the expected fraction of pig to one sixteenth, and so forth. The porcine fraction of the genome would therefore rapidly decrease, so that in just a few generations it would become much harder to detect.

If the same process is considered from the standpoint of percent nucleotide identity with bonobos, the F_1_ genome would be around 90% similar (the pig half of an F_1_ hybrid’s genome would be about 80% identical to a bonobo’s, and the bonobo half, 100% identical). The second generation—the B_1_ generation produced by backcrossing to bonobos—would be about 95% identical to bonobos. In the B_2_ generation, the identity to bonobo would rise to approximately 97.5%, and so forth. (Note that 80%, the figure used here, does not represent the exact difference between bonobo and pig. Nor does it need to. With backcrossing, the numbers would quickly approach 100%, even if the initial number were much smaller, say 60%, in which case the series would be 80, 90, 95, 97.5, etc.)

Conversely, the percentage of nucleotides that would be identical to pig but different from bonobo would decline rapidly with each generation: 10%, 5%, 2.5%, and so forth. And, obviously, as the percentage of porcine DNA went down, it would become increasingly difficult to detect.

Still, if the process stopped here and any DNA of porcine origin in the hybrids remained restricted to a few large blocks, it would be a straightforward matter to find it with conventional techniques. But any population that arose by such means would at some point have to cease mating with bonobos and instead begin to breed mostly *inter se*. Otherwise, it would suffer the usual fate of any small hybrid population that repeatedly backcrosses to a large parental population. In only a few generations, the descendants of the original cross would become nearly indistinguishable from bonobos.

This sexual breakaway from the bonobo parental population would mark the outset of a new period of breeding in isolation, lasting right up to the present day, during which there would be further masking of any porcine vestiges at the nucleotide level. Throughout the many generations of this second stage, any blocks of porcine DNA still present after the first few generations of backcrossing would be thoroughly shredded and reassembled by meiotic recombination. This stage would begin with a population of individuals whose genomes were composed mostly of large blocks of bonobo origin but also of a lesser fraction of large porcine blocks. The bonobo and porcine blocks would differ in their relative proportions and be arranged differently in different individuals. When the members of this varying population interbred, these structural differences in their chromosomes would result in unstable meiosis in the gonads of their offspring. So, in every new generation, there would be many new chromosomal inversions, translocations, losses, duplications and breakages. The result would be a shortening of the average block size over time as well as a restructuring of the individual chromosomes. Any traces of pig would gradually become a sprinkling of relatively short fragments.

With time, this blending of bonobo and porcine elements would become exceedingly intimate, given that during meiosis there would also be, in every generation, exchanges of strands of DNA between chromosomes, a process known as “crossing-over.” In the long run, crossing-over would have an effect similar to that of a meat grinder. If one took a mixture of 97.5% beef and 2.5% turkey and ran it through a grinder a thousand times, the turkey would become much harder to detect than if it were left as a single, uniform mass embedded in a pile of ground beef. It would have been chopped up into tiny pieces and intimately mixed with the beef. In the same way, after thousands of rounds of meiosis, it would be much harder to find any short stretches of DNA in humans that contained nucleotides that both matched pig and differed from bonobo.

Strands of DNA would not only be exchanged during each of these repeated rounds of meiosis but would also be altered with respect to their sequence content. Since DNA is double-stranded, when a single strand from one chromosome replaces a single strand on another during crossing-over in meiosis I, there can be mismatched nucleotides (in a nucleotide base pair, *A* must always pair with *T*, and *G* always with *C*). The cell’s mismatch repair machinery resolves any such incompatible pairs by replacing one of the pair’s nucleotides with one that matches the remaining one (Manhart and Alani 2015). Therefore, in the specific case now under discussion, pig-matching nucleotides would sometimes be replaced with bonobo-matching nucleotides and sometimes vice-versa. Since these replacements would happen in different ways in different individuals, different people would match pig at different nucleotide positions, which with passing generations, might be expected to result in an oscillation at such positions between pig- and bonobo-matching nucleotides that continued indefinitely.

If these things really happened, any human genome would be composed of two components, a large one made up of bonobo-matching nucleotides, and a smaller one composed of ones that did not match bonobo, but matched pig instead. *Moreover, because the nucleotides in the human genome that did not match bonobo would have been put in place by a process that replaced bonobo DNA with pig DNA, the fraction of human nucleotides that matched pig and did not match bonobo would be essentially the same as the fraction of nucleotides differing from bonobo*. The results of this study show that such is in fact the case. Such a finding is surely not expected under the divergence hypothesis (as the Discussion section will explain).

## Methods

In this study, a new data miner, BOOMSTICK, was used to search the human and bonobo genomes for matches to a large, random selection of short porcine nucleotide sequence queries.

### Queries and matches

BOOMSTICK searches one genome, the *target genome*, for matches to queries from a second genome, the *query genome*. Before the searches begin, hundreds of thousands of queries are stored in a single file, which will be called the *query file*. BOOMSTICK takes all the queries in the query file and searches a nucleotide sequence file from the target genome, the *target file*, for matches.

The name of the program was chosen because *boomstick* is slang for a sawed-off shotgun, and the program implements an algorithm that resembles a shotgun blast: a salvo of queries, analogous to the many pellets fired by a shotgun, is directed against the target file. The target genome is therefore searched, not for matches to a single query, as is the case in ordinary BLAST, but instead for matches to a vast number of different queries.

BOOMSTICK will accept queries of varying length. A query length of forty nucleotides was chosen, and a “match” defined as any sequence in the target genome that matched one of these 40-mer queries at a minimum of 35 positions. This criterion was selected because it (1) would allow detection of sequences that had been obscured by mismatch repair or random mutations but (2) would at the same time be true, since the probability of obtaining such a match at random is vanishingly small. The probability of a 40-mer query being identical at 35 or more positions to a randomly generated series of 40 *A*s, *G*s, *C*s, and *T*s is 3.8 × 10^-61^ (per the University of Michigan Statistics Online Computational Resource high precision binomial distribution calculator; https://tinyurl.com/juyj5se4). This very low probability of such a match occurring at random makes it nearly certain that the nucleotide positions being compared are equivalent (i.e., that they share common ancestry). And yet 40-mer queries are short enough to work well with the alignment algorithm used by BOOMSTICK, which does not allow for gaps.

BOOMSTICK was used to scan two target genomes, one representing *Homo sapiens*, the other, *Pan paniscus* (bonobo) with queries from a *Sus scrofa* (domestic pig) query genome. To do so, a query file was first created containing 813,194 porcine 40-mers sampled at 3,000-nucleotide intervals along each of the 20 pig chromosomes (18 autosomes and two sex chromosomes). A 3,000-nucleotide interval was used because it allowed batches of jobs to complete within a convenient 24-hour cycle. Since there is no reason to believe that a periodicity exists in the pig genome, this method of selecting queries yields a random sample. All porcine query sequences used in this study were sampled from the source indicated in Table 1. The specific breed of pig in question was Duroc. The same query file was used in all scans. For reasons that will be explained, a chimpanzee Y-chromosome was searched as well.

**Table 1.**
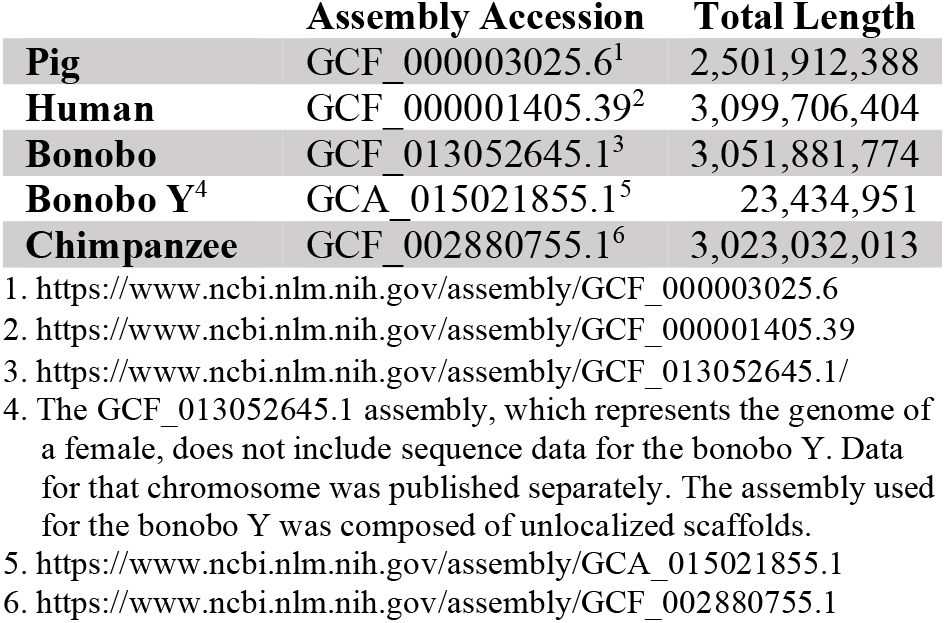
Sources of the nucleotide sequence files used

**Table 2.**
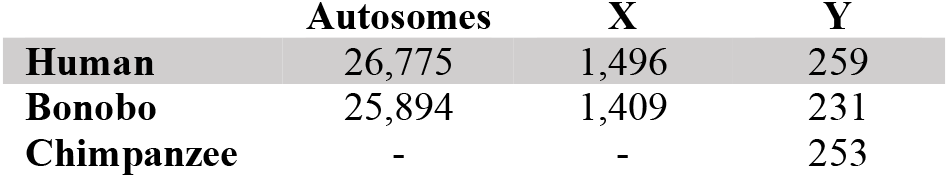
Number of euchromatic segments scanned.

|  | **Autosomes** | **X** | **Y** |
| --- | --- | --- | --- |
| **Human** | 26,775 | 1,496 | 259 |
| **Bonobo** | 25,894 | 1,409 | 231 |
| **Chimpanzee** | - | - | 253 |

### Scans limited to euchromatin

The only respect in which the query sample departed from being random was in that parameter settings were used that allowed BOOMSTICK to scan only single-copy (euchromatic) sequences. The settings activated filters that reject simple tandem repeats (e.g., mononucleotide, dinucleotide and trinucleotide repeats). So, the program did not seek matches to such repeats. Activation of these filters did not, however, bias the results, because the same set of queries and the same parameters were used to process all the nucleotide sequence target files used to represent the human and bonobo genomes. Euchromatic genes are nearly all transcriptionally active. In the human genome euchromatin comprises about 90% of the DNA.

### Segments

The target files used in the scans were extracted from the sources indicated in Table 1. For each of the three target genomes, the UNIX split command was used to divide the nucleotide sequence in each chromosome file into smaller 100-kB files (102,400 bytes), each containing a different segment of the chromosome. Each segment was 101,136 nucleotides long (the GREP command-line utility was used to verify that the 100-kB segment files contain 102,400 characters, of which 101,136 represent nucleotides and 1264 are end-of-line characters). These segments were contiguous and non-overlapping. Hence, they represent each chromosome in its entirety, except for regions composed of constitutive heterochromatin, which were not investigated in this study. The original chromosome files were parsed in this way for two reasons. One was to facilitate rapid parallel processing. The other was to make the human and bonobo samples comparable.

For each parsed chromosome, there was a terminal file generated which was smaller than 100 kB. No results are reported for these remainder segments, and they were not included in the calculations of mean match rates, given that they are not only shorter than the other, full-length segments, but also vary widely in length, so that their match counts are not comparable.

The sequence file for the bonobo Y-chromosome contained only unlocalized scaffolds. To make it scannable by BOOMSTICK the FASTA headers were stripped out; the residual 3,590 individual sequences remaining were concatenated, and the UNIX split command was then used, as usual, to parse the resulting sequence into 100 kB target segment files. Given the unassembled status of the bonobo Y, results from scanning a chimpanzee Y are reported as well.

### Scans

BOOMSTICK was used to scan each of a total of 52,669 100-kB single-copy (euchromatic) autosomal target segment files for matches to each of the 813,194 porcine 40-mers in the query file, a total of 42,830,114,786 searches. It was also used to scan 2,906 100-kB sex-chromosome segments with the same query file, 2,363,141,764 additional scans. A UNIX shell script was used to submit each segment to a separate processor of the University of Colorado’s Summit supercomputing cluster (Anderson et al. 2017). Typically, a thousand separate segments were searched in a thousand simultaneous runs.

BOOMSTICK is programmed so that, in each segment file, no more than a single match can be found to each query, and any match already found to one query cannot be found as a match to any subsequent query. BOOMSTICK’s output includes the number of matches found in each segment. The match counts were used to calculate the mean number of matches per single-copy (euchromatic) segment for each study organism (see Results). In humans, approximately 90% of the DNA is euchromatic, as has been mentioned.

### Heterochromatic regions

Grunau et al. (2006) state that “In humans, regions surrounding centromeres and telomeres are heterochromatic, whereas large parts of the chromosome arms consist of transcriptionally competent euchromatin.” Heterochromatic regions, which compose about 10% of the human genome, are difficult to sequence because they are generally composed of repetitive DNA, which results in ambiguities during assembly. In this paper, results are reported only for single-copy regions (euchromatin), in part because heterochromatic regions often have only been partially sequenced, but mainly because, with the parameter settings used in this study, BOOMSTICK rejects repetitive queries, such as AAAAAAAAAA or TGTGTGTGTG. If such queries had been allowed, and if each of those thousands had potentially found thousands of matches in the target genome, run times would have mushroomed.

Submitted nucleotide sequence files often either omit heterochromatic regions or represent them by long strings of *N*s, a process known as “hard masking” (in a nucleotide sequence file, an *N* serves a place-holder function, in that it merely indicates a position in the sequence for which the nucleotide’s identity could not be determined). In preparation for searching a segment, BOOMSTICK first loads into memory the nucleotide sequence in a target segment file. It then preprocesses the sequence to ascertain the number of *N*s that it contains. When more than half the nucleotide positions in the segment are occupied by *N*s, processing of the segment aborts. Such segments may be termed “N-rich.”

And, as stated above, with the parameter settings used in the searches, BOOMSTICK also rejects repetitive queries. Therefore, segments that are not *N*-rich but do contain only repetitive sequence cannot generate matches with BOOMSTICK as it is currently programmed. Such segments, generating no matches, but that are not *N*-rich, may be called “zero-hit.”

*N*-rich segments do not enter into the analyses presented in this paper, since BOOMSTICK rejects them without ever actually searching them for matches to queries. Zero-hit segments, however, pose a special problem because it is not immediately clear, from the simple fact that they have failed to generate matches, whether they are heterochromatic or euchromatic. In the former case, their failure to produce matches would be due to BOOMSTICK’s rejection of repetitive queries. But in the latter, they might represent segments that truly contained no matches and that therefore should be included in the calculations of mean segmental match rates (MSMRs). It seemed unlikely that zero-hit segments were of the former type (i.e., heterochromatic), as are *N*-rich segments, given that it seemed unlikely that a scan of 100,000 nucleotides with 814,194 randomly selected 40-mers would generate no hits at all. But it seemed prudent to check whether this surmise was correct.

A cursory analysis of the results for human chromosome 1 revealed that both *N*-rich and zero-hit segments for that chromosome did fall into highly repetitive regions—the centromere and a large pericentromeric region, both of which are known to be composed of constitutive heterochromatin. To further ascertain whether such segments represent heterochromatin, a detailed evaluation of the human Y chromosome was conducted. For that chromosome, BOOMSTICK was used to evaluate 566 segments. Of these, 259 were neither *N*-rich nor zero-hit, 301 were *N*-rich, and six, zero-hit. To show that the *N*-rich and zero-hit segments from the human Y do correspond to repetitive regions, the position of each such segment on the Y was calculated and NCBI’s Genome Sequence Viewer was used to compare those calculated positions with the known positions of heterochromatin on that chromosome. Such segments, which on the human Y occur in four separate regions composed of contiguous *N*-rich or zero-hit segments, were in fact found to occur in high-repeat regions and not elsewhere (see Table 3). Therefore, on the human Y, *N*-rich segments represent regions that had, for the most part, not yet been sequenced in the assembly scanned, and zero-hit segments represent regions that had in fact been mostly sequenced, but that were composed of heterochromatin. The close agreement between the calculated positions of the *N*-rich and zero-hit segments found for the human Y to the known positions of heterochromatic regions on that chromosome allayed any doubts as to whether such segments were in fact heterochromatic. They have therefore been omitted from the analysis, not only for the human Y but also for all other chromosomes.

**Table 3.** Calculated positions of *N-*rich and zero-hit segments compared to the positions of heterochromatic regions on human chromosome Y (positions are given in megabase pairs from the distal end of the p arm).

| **Positions<sup>‡</sup> of *N*-rich**<br>**and zero-hit segments**<br>**(per BOOMSTICK)** | **Positions of**<br>**heterochromatin (NCBI**<br>**Genome Data Viewer )** |
| --- | --- |
| 0.0 - 0.20 | 0.0 - 0.22* |
| 10.31 - 11.73 | 10.27 - 11.03<sup>†</sup> |
| 20.12 - 20.33 | 20.20 - 20.23 |
| 26.7 - 56.8 | 26.3 - 56.8<sup>‡</sup> |

\*. This region corresponds to the p-arm telomere.
<sup>†</sup>. This region corresponds to the centromere.
<sup>‡</sup>. This region corresponds to the NRY region.
‡. When each chromosome was parsed into a set of contiguous, non-overlapping segments, each segment file was assigned a number corresponding to its ordered position along the chromosome, starting with segment 1, taken from the extreme distal end of the p arm. Since each segment file corresponds to 101,136 nucleotides, the position of the end of each *N*-rich and zero-hit segment is simply its segment number multiplied by 101,136. The position of its beginning can be calculated by subtracting 101,135 from the position of its end.

## Results

For the set of 813,194 randomly selected porcine 40-mer queries used in this study, BOOMSTICK found more matching sequences in the autosomes and Y chromosome of humans than in those of bonobos. When scanned with the same query set, the match rate of the human X chromosome did not differ significantly from that of bonobos. In all cases, only single-copy (euchromatic) regions were searched. *The key finding was that essentially all human autosomal euchromatic nucleotides that differ from the bonobo nucleotide at the corresponding position, match pig*.

### MSMRS

From the data produced by the scans, the mean number of matches per segment—what is here called the mean segmental match rate (MSMR)—was calculated for both humans and bonobos. This was done for the single-copy (euchromatic) portion of each autosome and of each of the sex chromosomes. A higher MSMR indicates more matches to pig queries were found.

#### Autosomes

An autosome is any chromosome other than a sex chromosome. In a woman’s genome, they contain about 90% of the DNA, in a man’s, about 95%. In both humans and bonobos, autosomes are present in pairs. The MSMRs for the euchromatic regions of the autosomal genomes of both humans and bonobos are shown in Table 5. The human MSMR was 1.331% higher than the bonobo MSMR (Table 6).

**Table 5.** Autosomal MSMRs.

|  | Segments | Matches | MSMR |
| --- | --- | --- | --- |
| **Human** | 26,775 | 1,481,576 | 55.334 |
| **Bonobo** | 25,894 | 1,414,017 | 54.607 |

**Table 6.** Ratio between autosomal MSMRs.

|  | Ratio | Percent increase |
| --- | --- | --- |
| **Human/bonobo** | 1.01331 | 1.331% |

**Table 7.** X-chromosome MSMRs.

|  | Segments | Matches | MSMR |
| --- | --- | --- | --- |
| **Human** | 1,496 | 141,467 | 94.56 |
| **Bonobo** | 1,410 | 134,273 | 95.23 |

#### X chromosomes

Calculated MSMRs did not differ significantly between the human and bonobo X chromosomes. The MSMRs for the X chromosomes were the highest for any chromosomes included in this study, presumably because X chromosomes are strongly conserved among mammals, which would tend to increase the number of pig queries found on bonobo and human Xs. It has long been recognized that humans are more similar to panins with respect to the X chromosome than with respect to the autosomes.

#### Y chromosomes

Table 8 indicates the MSMR not only for the euchromatic fraction of a human and a bonobo Y chromosome, but also for a chimpanzee Y. Table 9 gives the ratios between the human Y-chromosome’s MSMR and that of each of the two panins. Note that the human Y-chromosome MSMR is considerably higher than that of either panin. The chimpanzee MSMR (57.24), however, seems the more reliable of the two panin MSMRs, given that the chimpanzee Y sequence was fully assembled, whereas the bonobo assembly was made up solely of unlocalized scaffolds. The increase in MSMR for humans versus bonobos is more marked for the Y than for any other chromosome.

**Table 8.** Y-chromosome MSMRs.

|  | Segments | Matches | MSMR |
| --- | --- | --- | --- |
| **Human** | 259 | 16,666 | 64.35 |
| **Bonobo** | 231 | 11,043 | 47.81 |
| **Chimpanzee** | 253 | 14,482 | 57.24 |

**Table 9.** Ratios between Y-chromosome MSMRs.

|  | Ratio | Percent increase |
| --- | --- | --- |
| **Human/bonobo** | 1.346 | 34.6% |
| **Human/chimpanzee** | 1.124 | 12.4% |

### Pig-Matching Nucleotides (PMNS)

In this study, a “match” was defined as any 40-mer identical to one of the porcine queries at 35 or more positions (see Methods), and each such position in a match, at which the pig query nucleotide is identical to the corresponding target nucleotide, is called a *pig-matching nucleotide* (PMN).

#### Autosomes

The total number of PMNs found by BOOMSTICK in the euchromatic regions of human and bonobo autosomes were calculated. To do so, the log files for the scans were parsed to determine the number (35, 36, 37, 38, 39 or 40) of identical positions for each match found by BOOMSTICK. All matches, for each target organism, more than a million matches each, were classified in this way. For both humans and bonobos, the total number of PMNs, as well as the mean number of PMNs per segment, were then calculated. These totals and the ratios between them are shown in tables 10 and 11. Note that the ratios and percentages based on PMNs are the same as those calculated from MSMRs.

**Table 10.** PMNs per autosomal segment.

|  | Segments scanned | Pig-matching nucleotides |  |
| --- | --- | --- | --- |
|  |  | Total | Segmental Mean |
| **Human** | 26,775 | 52,307,723 | 1953.60 |
| **Bonobo** | 25,894 | 49,924,271 | 1928.02 |

**Table 11.** Ratio between the mean autosomal PMNs.

|  | Ratio | Percent increase |
| --- | --- | --- |
| **Human/bonobo** | 1.01327 | 1.327% |

#### Human autosomes

When the 1,481,843 human euchromatic autosomal matches were classified, 1,132,699 were identical at 35 positions, 274,328 at 36, 54,422 at 37, 14,153 at 38, 4,191 at 39, and 2,050 at 40.

Thus, the mean number of PMNs per query match was 35.305461 = [(1,132,699 × 35) + (274,328 × 36) +

(54,422 × 37) + (14,153 × 38) + (4,191 × 39) + (2,050 × 40)]/1,481,843. This figure was calculated using all matches found, including those in the 22 remainder segments generated when the 22 human autosomes were parsed by the file-splitting utility (see Methods). It can therefore be calculated that in humans there were 52,307,723 (= 35.305461 × 1,481,576) PMNs in the 1,481,576 matches BOOMSTICK found in full-length (non-remainder) autosomal segments. There were 26,775 such segments. Hence, the mean number of PMNs found per segment in humans was 1,953.60 (= 52,307,723/26,775).

#### Bonobo autosomes

When the 1,414,497 bonobo euchromatic autosomal matches were classified, 1,081,148 were the same at 35 positions, 260,844 at 36, 52,705 at 37, 13,669 at 38, 4,093 at 39, and 2,038 at 40.

Thus, the mean number of PMNs per match was 35.306698 = [(1,081,148 × 35) + (260,844 × 36) + (52,705 × 37) + (13,669 × 38) + (4,093 × 39) + (2,038 × 40)]/1,414,497. Again, this figure was calculated from all autosomal matches, including those from the 23 bonobo remainder segments. There would therefore be 49,924,271 (= 35.306698 × 1,414,017) PMNs in the 1,414,017 matches BOOMSTICK found in full-length bonobo autosomal segments. But there were 25,894 such segments. So, the mean number of PMNs per segment was 1,928.02 (= 49,924,271/25,894).

#### Autosomal ratios

So, if humans are compared with bonobos by calculating the ratios of the mean number of PMNs found per euchromatic autosomal segment, humans had 1.01327 (= 1953.60/1928.02) times as many as did bonobos, an increase of 1.327% (very nearly the exact ratio calculated from MSMRs).

## Discussion

The Introduction explained that the hybrid hypothesis of human origins is the idea that humans have descended from one or more ancient hybridization events involving bonobo and pig, the genetic traces of which would have been much obscured by subsequent multiple generations of backcrossing to bonobos, followed by thousands of generations of masking meiotic recombination within the human lineage. Therefore, if the hybrid hypothesis is correct, the human nucleotides differing from bonobo should largely match pig. Conversely, if humans and bonobos have simply diverged from a common ancestor, there is no expectation that those human nucleotides differing from those of bonobos should all match pig.

But the results of this study are consistent with the former of these two possibilities. They show that the excess of pig-matching nucleotides (PMNs) in the human autosomal genome closely corresponds with the reported percent nucleotide difference between the human and bonobo autosomal genomes. Indeed, the findings indicate that essentially *all* of the human euchromatic autosomal nucleotides that differ from bonobo euchromatic autosomal nucleotides are pig matching nucleotides, as will now be explained.

Humans and bonobos are diploid organisms, so they have two copies of each autosome and therefore two copies of each autosomal gene. These are sometimes differing copies, that is, alleles. Prüfer et al. (2012) assembled the genome of a female bonobo named Ulindi and compared it to those of humans. According to those authors, a randomly selected human autosomal gene differs from its bonobo counterpart, on average, at 1.3 nucleotide positions in a hundred, that is, at a rate of 1.3%.

It is remarkable, then, that in the present study found that the number of pig-matching nucleotides detected by such a large number (813,194) randomly selected 40-mer pig queries was 1.327% higher in human autosomes than in bonobo autosomes, that is, almost exactly 1.3%. So, the number of nucleotide positions that are identical to pig in human autosomes but different in those of bonobo, is substantially identical to the number of nucleotide positions that Prüfer et al. reported to differ between humans and bonobos. *This means that essentially all the autosomal nucleotides differentiating humans from bonobos are pig-matching—a result completely unexpected under the divergence hypothesis*.

The hybrid hypothesis does account for the identity of these two numbers. Under that view, the genetic differences between humans and bonobos arose through a process that added pig DNA to the bonobo genome. Therefore, the genomic process that created the differences between humans and bonobos was the same process that produced an excess of pig-matching nucleotides in humans, as compared to bonobos.

In contrast, under the divergence hypothesis, humans and bonobos share a recent common ancestor (∼5 MYA), whereas humans and bonobos share with pigs only a very ancient one (∼60 MYA). With such assumptions, it is hard to see why all the nucleotides distinguishing *Homo* from *Pan* would all match *Sus*. Imagine a series of mutations affecting the bonobo and human lineages during their divergence from their common ancestor, as described in the divergence hypothesis. And suppose that in the end this process produced two genomes, one for modern humans and one for bonobos. Under such circumstances, why would all the nucleotide positions that differed in human and bonobo, match pig in humans? There is no reason to expect any such bias, let alone a bias so completely lopsided. But in fact, essentially all such differentiating nucleotides in human autosomes do match pig. The hybrid hypothesis, then, is at least provisionally supported by the findings of this study, whereas the divergence hypothesis is not.

Note that thinking in terms of the divergence hypothesis would lead one to deem organisms with more human-matching nucleotides (HMNs) more closely related to humans than those with fewer. And many organisms do have more HMNs than do pigs, for example, any non-human primate. It must be emphasized, however, that counts of HMNs cannot in themselves be used to discriminate between the divergence and hybrid hypotheses.

Indeed, even certain non-primates might well have higher hit rates than pig against humans and panins. Even under the hybrid hypothesis there is no reason to suppose that humans would necessarily have a higher level of nucleotide identity with pigs than with any other non-primate. By the fact that genetic variation exists among the various kinds of non-primates, we know that some animals in that category will have a higher level of nucleotide identity with bonobos than will others. And those will have a higher level of identity with humans as well, because the genomes of humans and bonobos are identical at 98.7% of the positions they contain. One can therefore easily imagine that there would be certain non-primates that shared a higher level of nucleotide identity with humans and bonobos than pigs do, even if they themselves did not participate in the cross.

The finding that the excess of pig-matching autosomal nucleotides in humans (versus bonobos) equals the number of autosomal nucleotides by which humans differ from bonobos, however, remains a striking, remarkable and unexpected result, which is clearly inconsistent with the divergence hypothesis.

### Structural rearrangement

But it must be considered, too, that the findings of this study were not produced in a vacuum. There is a separate line of genomic evidence—the existence of structural differences distinguishing human chromosomes from those of panins, differences known as *structural rearrangements*. Comparison of humans with panins has revealed many differences with respect to the presence/absence of blocks of DNA, as well as numerous inversions. Kronenberg et al. (2018) searched for regions fifty base pairs or more in length that, in comparison with panins, have either been added to or deleted from the human genome. They counted 11,897 insertions and 5,892 deletions. Kehrer-Sawatzki and Cooper (2007) say, “Genome-wide comparisons have indicated that some 40-45 Mb of lineage-specific sequence results from insertions/deletions resulting in copy number differences between *HSA* and *PTR*” (i.e., human and chimpanzee).

If humans are the descendants of ancient hybridization between bonobos and pigs, it is easy to understand why the human and bonobo genomes differ in this way, with respect to many insertions, deletions and inversions. When the chromosomes of the parents in a hybrid cross differ with respect to structural rearrangements—as do pigs and panins—meiosis becomes chaotic so that blocks of DNA in the genomes of their hybrid descendants will be frequently inverted, deleted, duplicated and transposed (Grant 1985, McCarthy et al. 1995; Stebbins 1957, 1958). In short, structural differences between the participants in a hybrid cross result in the production of new structural rearrangements in their hybrid descendants. Some of these many rearrangements can eventually become established and come to characterize the stabilized hybrid derivative (McCarthy et al. 1995). Under such circumstances there is selection for fuller structural homozygosity, that is, for better chromosome pairing, which is correlated with higher levels of fertility (ibid.).

One can think of the process as a feedback loop converging on a stability point: (1) Individuals with higher pairing produce more offspring; (2) those offspring inherit higher pairing; (3) any of those offspring with higher pairing produce even more offspring, with even higher pairing. So, the general level of pairing rises with each cycle and the process converges on a state where full pairing is achieved. In stochastic spatial computer simulations of this process (McCarthy et al. 1995), hybrid populations that began with high levels of structural heterozygosity consistently, and seemingly inevitably, transitioned to stable populations composed of individuals with fully paired identical chromosome sets distinct from those of the parents participating in the original F_1_ cross.

In the same way, in a situation where bonobos crossed with pigs to produce a variable hybrid swarm, a preponderance of the resulting hybrids would be structural heterozygotes with many improperly paired chromosomes. But the process described in the previous paragraph would produce a new set of much rearranged, but fully paired chromosomes, different from those of pigs and panins, and characteristic of human beings.

In contrast, there seems to be no satisfactory explanation for the existence of these rearrangements in terms of the divergence hypothesis. Under that view, one can understand how, say, an inversion might arise *de novo*, as a gross mutation. However, chromosomes bearing such rearrangements tend not to spread through an otherwise uniform population, because inversion heterozygotes are at a severe reproductive disadvantage, as poor chromosome pairing means sub-par gamete production (Kauppi et al. 2012). And yet, not one but many inversions differentiate the human genome from those of panins, which raises the question of how these large-scale structural differences between the human and panin genomes have become established.

### Y chromosome

Under the hybrid hypothesis, the greater similarity of *Homo* to *Pan*, than to *Sus*, suggests that backcrossing occurred to bonobo. Mammalian hybrids upon reaching sexual maturity tend to choose mates of their mother’s kind. This bias is due to imprinting, the biological phenomenon in which adults prefer to mate with individuals of the same kind as those who raised them (Lorenz 1952). Backcrossing to bonobo, then, would be more likely if the mother in the F_1_ cross had been a bonobo. Otherwise, the F_1_ hybrids would have been raised by a sow and therefore have imprinted on pigs and have sought porcine mates. All of which suggests the F_1_ cross between bonobo and pig would have involved a female bonobo and a male pig. Ethological and anatomical considerations also suggest that a female panin would be more likely to mate with a boar than a male panin with a sow (McCarthy 2008b).

Therefore, under the hybrid hypothesis, it isn’t surprising that the X chromosomes are more similar in bonobos and humans than are the autosomes. Previous studies have shown that the chimpanzee and bonobo Xs are more than 99% identical to those of human beings (Kehrer-Sawatzki and Cooper 2007; Patterson et al. 2006), which agrees well with this study’s finding that the human and panin Xs do not differ significantly when scanned for matches to pig queries. In the scenario described in the previous paragraph, the bonobo mother would pass a bonobo X to the F_1_ hybrid(s), whereas the boar father would pass either a pig X or a pig Y. Thus, gametes produced by an F_1_ hybrid could contain a bonobo X, and either a pig X or a pig Y. Given that the bonobo and human Xs are so similar, it seems evident that no pig X could have passed to the descendant backcross hybrids. The remaining two possibilities are that the backcross hybrids had either two bonobo Xs (one from the F_1_ parent and one from the bonobo backcross parent) or a single bonobo X (from the bonobo parent) and a pig Y (from the F_1_ parent).

In this way, the initial backcross (B_1_) hybrids could acquire an intact bonobo X. With subsequent generations of backcrossing to bonobo, the presence of a bonobo X in any later-generation hybrids would be virtually assured. But the passage of at least some genetic material from pig autosomes to later generations would have been inevitable. Through long-term, repeated recombination with bonobo autosomes, this passed porcine DNA, and the shredded fragments descended from it, would surely alter the original bonobo autosomes to be more like pig. The fact that this study’s results show that pig-matching nucleotides in the human Y chromosome were far more numerous than in bonobo Ys is consistent with previous reports stating that the human and panin Ys are radically different. Hughes et al. (2010) remark that with respect to all chromosomes other than the Y, chimpanzees and humans are quite similar. But, regarding the male specific region of the Y chromosome (MSY), they say that “at six million years of separation, the difference in MSY gene content in chimpanzee and human is more comparable to the difference in autosomal gene content in chicken and human, at 310 million years of separation.” This extreme dichotomy between the Ys of two organisms that are so similar with respect to their other chromosomes is hard to understand under the divergence hypothesis.

Under the hybrid hypothesis, however, this glaring difference between the human and panin Ys is easily explicable. Under that view, some, as yet unidentified, elements of the human Y would be derived from the pig progenitor. The greater persistence of pig-matching nucleotides in the Y would be due to the Y’s existing as a single copy and mostly lacking a recombination partner. As Bradbury (2017) points out, “The majority of the [human] Y chromosome (95% of the Y chromosome) contains DNA that does not undergo recombination.” Any components of the Y acquired early on from the porcine forebear would therefore be buffered against the masking effects of initial backcrossing and the subsequent thousands of generations of meiotic recombination (see Introduction). The fact that the human Y has been extensively restructured versus the panin Y (Hughes et al. 2010), would be the result of the many rounds of aberrant meiosis that would be expected during gametogenesis in a hypervariable population of bonobo-pig hybrids. Such atypical recombination may have been amplified by the occurrence of individuals with multiple Ys (XYY individuals occur at appreciable frequencies in the human population even today). Ishishita et al. (2015) reported abnormal pairing of autosomes with X and Y sex chromosomes during meiosis I in hamster hybrids.

### Glans penis and clitoridis

Y chromosomes are, of course, found only in males. Interestingly, men have at least one feature that distinguishes them from both women and male panins, and which may have had its origin in the hypothesized cross: a glans penis. The panine glans clitoridis is almost identical in form to the human glans penis, but male panins have no true glans (Ashley-Montagu 1937; Dixson 1987, p. 429, Fig. 3h; Hill 1972, p. 115; Izor et al. 1981; Martin and Gould 1981; Savage-Rumbaugh and Wilkerson 1978, Plate 4; Sonntag 1924, p. 270; de Waal and Lanting 1997, p. 127). Nor does a glans occur in boars. These facts suggest that genes governing the development of this structure have somehow been transferred from the bonobo X to the human Y. Under the hybrid hypothesis, such a transposition would have occurred during the aberrant meiotic recombination that would characterize a structurally heterozygotic population of bonobo-pig hybrids, so that some genes in the bonobo X were incorporated into what became the human Y. The traits common to the human penis and panine clitoris would then be specified in men by these transferred genes present in the human MSY. And indeed, Hughes et al. (2010; citing Page et al. 1984) say that one sequence class “in the human MSY euchromatin—the X-transposed sequences—has no counterpart in the chimpanzee MSY. The presence of these sequences in the human MSY is the result of an X-to-Y transposition that occurred in the human lineage after its divergence from the chimpanzee lineage.” They state (ibid.) that the genes in question are *TGIF2LY* and *PCDH11Y*. The former of these two, *TGIF2LY*, encodes a member of the *TALE*/*TGIF* homeobox family of transcription factors. Homeobox genes are involved in the regulation of patterns of anatomical development. There is therefore reason to suspect that *TGIF2LY* functions in the development both of the human glans penis and the panine glans clitoridis.

### Phenotypic traits

Even before the launch of this survey with BOOMSTICK, a supporting, independent line of evidence—the many porcine traits distinguishing humans from bonobos—was arresting enough to make a genetic analysis seem worthwhile. A partial list of such traits, as documented by McCarthy (2008b), nearly all of which distinguish humans not only from bonobos, but also from all nonhuman primates, includes the following.

Sparse pelage (i.e., largely naked skin)

Piglike, cartilaginous snout

Ubiquitous layer of subcutaneous fat (panniculus adiposus)

Cutaneous muscle present only in neck and face

Melanocytes present in the matrix of the hair follicles

Darwin’s point

Musculocutaneous arteries

Atherosclerosis

Lightly pigmented irides (absent in nonhuman primates)

White, visible sclera surrounding iris

Diverticulum in fetal stomach

Vocal ligaments

Epidermal lipids

Cranial emissary foramina

Pharyngeal and nasal blood plexuses

Multipyramidal kidneys

Prostate encircling urethra

Bulbourethral glands

White of eye (absent in nonhuman primates)

Valves of Kerckring

Absence of ischial callosities

Absence of laryngeal sacs

Absence of os penis (baculum)

Absence of conspicuous sexual swellings in women

Indeed, it is widely known that porcine heart valves are used as replacements for human valves, and that pig skin is used in treating burn patients, as pig cutaneous anatomy is so similar to that of human skin. Pig kidneys look almost identical to those of humans, inside and out, while the renal anatomy of panins is markedly different from that of humans, both externally, and especially, internally.

As was pointed out in the Introduction, if a substantial number of these human traits, not found in other primates, were also not found in pigs, then one could reasonably attribute those that were in fact found in pigs to convergent evolution. But such is not the case. Instead, such traits are consistently found in pigs. This finding supports the hybrid hypothesis and was an original impetus for the present study.

### PHUNs

Call a human pig-matching nucleotide that does not match the equivalent position in bonobos a pig-human unique nucleotide (PHUN). The 26,775 human euchromatic autosomal segments scanned in this study contain 2,706,916,400 (= 101,136 × 26,775) nucleotides. In those segments, according to the data produced by BOOMSTICK, there are 3.593 × 10^7^ (= 0.01327 × 2,706,916,400) PHUNs. In the approximately 1.1 × 10^6^ (= 22(101,136)/2) nucleotides in the 22 remainder segments, there would be an expectation of finding an additional 1.5 × 10^4^ (= 0.01327(1.1 × 10^6^)) PHUNs. So, there would be about 3.594 × 10^7^ (= 3.503 × 10^7^ + 0.0015 × 10^7^) PHUNs in the sequence files of human euchromatic autosomal origin scanned for this study. But only a *single* copy of each autosome was scanned. Autosomes exist in pairs. The number of PHUNs in the euchromatic portion of an actual human autosomal genome, which is *diploid*, would therefore be at least twice that number, or about 72 million (7.188 × 10^7^) nucleotides. This autosomal figure represents a lower-bound on the number of PHUNs present in the human genome, since it leaves out of account any additional PHUNs lurking in the X and Y chromosomes, or within autosomal heterochromatin.

The injection of 72 million PHUNs would be expected to alter a bonobo genome radically, plausibly enough to produce that of a human being. A single copy of human chromosome 21, which contains 4.67 × 10^7^ nucleotides, amounts to only about 0.75% of a man’s genome and 0.73% of a woman’s. And yet, the addition of one extra copy of this chromosome (trisomy 21) is enough to produce Down syndrome, a condition that alters a wide variety of phenotypes. But adding a copy of a chromosome is a mere change in dosage and involves no change whatsoever in nucleotide sequence.

In humans, the dose of pig-matching nucleotides not found in bonobos is at least 1.5 (= 7.2/4.67) times as great as the dose of nucleotides distinguishing a person affected by Down syndrome from one unaffected. But the dose of PHUNs in the human genome is not the mere addition of an identical third copy of a chromosome, as in Down. Instead, it involves the replacement of more than 70 million nucleotides of bonobo DNA with pig-matching DNA. In short, the expected phenotypic effect would be profound. And presumably any such effect would, in men, be intensified, given the far greater frequency of pig-matching nucleotides seen in the human Y.

### Summary

The findings of this study are consistent with the hybrid hypothesis of human origins, the idea that the human lineage descends from ancient hybridization events involving bonobo and pig. Under that hypothesis,

- The human autosomes are, broadly speaking, extensively rearranged bonobo autosomes (altered by numerous insertions, transpositions, deletions and inversions) peppered with innumerable fragments of pig DNA.
- Within this chromosomal framework, due to the action of mismatch repair mechanisms during meiosis, there will be ongoing individual human variation with respect to the position of pig-matching nucleotides (see Introduction), with different people matching pig at different nucleotide positions, which would likely result in phenotypic variation.
- The human X chromosome is very close to the panin X, having been kept largely separate from porcine genetic influence
- Men are more affected than women by porcine ancestry because the human Y chromosome, though relatively small, is far more strongly influenced by pig ancestry than is any other human chromosome.
- For the first time, the huge phenotypic difference between humans and bonobos becomes understandable at the genetic level.

The divergence hypothesis, in which humans and bonobos share a recent common ancestor but only a very ancient one with pigs, is inconsistent with this study’s finding—that essentially all of the nucleotides distinguishing human from bonobo match pig. One needs no statistical test to see that under that view such a series of events could not happen at random. Nor does the divergence hypothesis explain the extensive rearrangement of human autosomes versus those of bonobos, nor the fact that the many human traits not seen in bonobos are nearly always found in pigs.

If the hybrid hypothesis continues to be supported, thinking in terms of it will suggest many new lines of inquiry in any field where the nature of human origins has an intellectual bearing.

## Acknowledgements

This work utilized the RMACC Summit supercomputer, which is supported by the National Science Foundation (awards ACI-1532235 and ACI-1532236), the University of Colorado Boulder, and Colorado State University. The Summit supercomputer is a joint effort of the University of Colorado Boulder and Colorado State University. Thanks are due, too, to Edward Falkowski for his assistance in submitting scans to the Summit supercomputer.

